# Development of the First Cytochrome Oxidase I Barcode and Evidence for a Single Haplotype Associated with the Recent United States Invasion of the Pasture Mealybug *Heliococcus summervillei* (Hemiptera: Pseudococcidae)

**DOI:** 10.64898/2026.08.04.742657

**Authors:** Muhammad Z. Ahmed, Peilin Tan, Nisha Yadav, Caroline Hauxwell, David R. Kerns, Blake Wilson, Nicole Quinn, Isaac L. Esquivel, Sachin Rustgi, Ynna Hernandez-Europa, Darcy Patrick

**Affiliations:** Department of Plant and Environmental Sciences, Pee Dee Research and Education Center, Clemson University, Florence, South Carolina, USA; School of Biology and Environmental Science, Queensland University of Technology, Brisbane, Queensland, Australia; Department of Entomology, Texas A&M University, College Station, Texas, USA; Department of Entomology, Louisiana State University Agricultural Center, Baton Rouge, Louisiana, USA; Institute of Food and Agricultural Sciences, Department of Entomology and Nematology, Indian River Research and Education Center, University of Florida, Fort Pierce, Florida, USA; Institute of Food and Agricultural Sciences, Department of Entomology and Nematology, North Florida Research and Education Center, University of Florida, Quincy, Florida, USA

**Keywords:** biosecurity, genetics, invasive pest, pest management, regulatory diagnostics, turfgrass

## Abstract

*Heliococcus summervillei* is an emerging invasive mealybug that causes severe dieback in grasses in pastures and turfgrass landscapes. It is widespread in Australia and has recently been detected across the Caribbean, Mexico, and the United States. Accurate mealybug identification is challenging because of cryptic and overlapping diagnostic characters and limited taxonomic expertise and literature, making molecular tools essential for regulatory diagnostics and management. We developed the first Cytochrome Oxidase I (COI) sequences for morphology-based Type A and Type B variants of *H. summervillei* and used them to examine mitochondrial variation across available populations. COI sequences reveal approximately a 10% mitochondrial split between the Type A and Type B variants. Phylogenetic, haplotype network, and genetic distance analyses show that all Type B populations share a single haplotype associated with a recent invasion of the United States and Australia. Type A in Barbados contains two closely related haplotypes that may represent a historically stable mitochondrial variant corresponding to the holotype’s morphological description. Together, these results establish the first COI reference library for *H. summervillei*, resolve mitochondrial lineage structure, and provide a practical tool for rapid identification and timely regulatory and pest management responses.

## 1 INTRODUCTION

Accurate identification of mealybugs (Hemiptera: Pseudococcidae) remains one of the most persistent challenges in biosecurity, regulatory diagnostics, quarantine decision-making, and pest management because many species are morphologically cryptic, highly polyphagous, and frequently intercepted at ports of entry (Ben-Dov 1994; Ahmed et al. 2015a,b; Ren et al. 2018; Ahmed et al. 2025a). Extreme intraspecific variation, overlapping and often cryptic morphological characters, and the requirement for highly technical and time-consuming slide-mounted preparations, together with limited taxonomic expertise and the continued reliance on adult-female-based keys, render mealybug identification slow and prone to error, particularly for immature stages (Williams & Granara de Willink 1992; Williams 2004; Malausa et al. 2011; Ahmed & Deeter 2022; Ahmed et al. 2023).

Deoxyribonucleic acid (DNA) barcoding using the mitochondrial cytochrome oxidase I (COI) has become the global standard for insect diagnostics due to its high amplification success, broad reference coverage in GenBank and the Barcode of Life Data Systems (BOLD), and robust potential for species-level resolution across Pseudococcidae (Hebert et al. 2003; Armstrong & Ball 2005; Park et al. 2011; Ratnasingham & Hebert 2013). For mealybugs specifically, COI is the most widely sequenced locus and forms the backbone of reference libraries used in regulatory diagnostics and pest management worldwide (Park et al. 2011; Ahmed et al. 2015a,b; Ren et al. 2018). Because most publicly available mealybug sequences correspond to COI, generating this sequence maximizes compatibility with existing databases and enables rapid, standardized identification.

Invasive species often show low genetic diversity during early establishment, but rapid population growth, host shifts, and insecticide exposure can generate genetic structure that affects diagnostics and management (Downie & Gullan 2004; De Barro & Ahmed 2011; Kumar et al. 2023; Al Naggar et al. 2025). Genetic variation within invasive populations can produce lineage-specific differences in biology (e.g., fecundity and thermal tolerance in *Bemisia tabaci* (Gennadius); Muñiz & Nombela 2001; Delatte et al. 2009; Park et al. 2021), dispersal potential (e.g., variable spread rates among *Thrips parvispinus* (Karny) haplotypes; Ahmed et al. 2025b), insecticide susceptibility (e.g., neonicotinoid-tolerant *Phenacoccus solenopsis* Tinsley lineages; Waqas et al. 2019; Shankarganesh et al. 2022), and ultimately the long-term sustainability of chemical control programs (e.g., pyrethroid-tolerant *Aleyrodes proletella* Linnaeus haplotypes; Springate & Colvin 2012) (Ahmed et al. 2015a; Xu et al. 2024; Ahmed et al. 2025b, 2026). Several mealybug complexes contain cryptic species or deeply divergent lineages that differ in host range (e.g., *Planococcus citri* (Risso) vs. *Pl. ficus* (Signoret); Downie & Gullan 2004; Malausa et al. 2011), developmental biology (e.g., lineage-level variation in *Phenacoccus* species and *Paracoccus marginatus* Williams & Granara de Willink; Malausa et al. 2011; Kumar et al. 2023), and compatibility with natural enemies (e.g., differential parasitism of *Ph. solenopsis* and *Ph. solani* Ferris by *Aenasius arizonensis* (Girault); Shankarganesh et al. 2022; Kumar et al. 2023).

These lineage-level differences influence integrated pest management (IPM) broadly, including but not limited to chemical control, host plant risk assessment, and biological control (De Barro & Ahmed 2011; Ahmed et al. 2015a; Kumar et al. 2023). For example, the success of classical biological control depends on matching parasitoids to the correct genetic lineage of the host (Meyerdirk et al. 2004; Muniappan et al. 2006; Ahmed et al. 2015a). Therefore, confirming genetic structure and identifying whether an invasion reflects a single introduction or multiple introductions using genetics is essential for risk assessment, spread prediction, and preventing further movement.

*Heliococcus summervillei* Brookes was first described as causing dieback in subtropical pastures in Australia in 1926 (Summerville 1928), whereas the species itself was formally described much later by Brookes (1978). More recently, it is an emerging invasive mealybug associated with severe pasture dieback in Australia (Schutze et al. 2019; Hauxwell et al. 2022a,b) and has recently been detected in the Caribbean, Mexico, and the continental United States (Gibbs 2020; Aca-Martinez et al. 2025; Biles et al. 2025; Powell & Hauxwell 2026; Wilson & Baca 2026). Although it has historically been reported from India, Pakistan, and New Caledonia (Brookes 1978; Brinon et al. 2004), its global genetic structure remains poorly understood. Recent work in Australia has revealed two distinct morphological variants: Type A (with morphology consistent with the holotype, Hernandez-Europa et al. 2026), which is thought to be the type from historical accounts of pest outbreaks and has become less prevalent over time, and Type B, a more invasive type that has become predominant in recent years. The types consistently differ in tibial pore structure and other diagnostic traits (Schutze et al. 2019; Hernandez-Europa et al. 2026; Powell & Hauxwell 2026). These differences raise the possibility of two species within *H. summervillei*, but no COI genetic data have been available to test this hypothesis.

Given the rapid spread of *H. summervillei* into the U.S. and the urgent need for reliable diagnostic tools, our first aim was to generate high-quality COI DNA sequences for *H. summervillei*. Because COI is the most widely sequenced mealybug marker in GenBank and BOLD, focusing on this region maximizes compatibility with existing databases and enables rapid regulatory actions. Our second aim was to analyze genetic variation across available invaded regions, including Australia, the Caribbean, and the U.S., to determine, with the data available at this stage, whether the U.S. outbreak represents a single introduction or multiple independent incursions. Third, we evaluated the potential for cryptic species or lineage divergence by comparing COI variation between the historical Type A and invasive Type B lineages. Finally, we used phylogenetic, genetic network, and genetic distance analyses to determine whether populations show genetic uniformity or detectable structure, providing insights relevant to chemical sustainability, biological control compatibility, and regulatory diagnostics.

## 2 MATERIALS AND METHODS

### 2.1 Sample collection

Specimens of *H. summervillei* were collected from multiple regions representing both historical and contemporary populations (Table S1). Barbados samples (NY73–NY77) were collected between 2020 and 2022 from pasture and lawn grasses. Additional North American samples included sugarcane from Louisiana (NY84), bermudagrass from Texas (NY85), limpograss from Florida (NY87–NY90), and centipede grass from the U.S. (NY99) (a new state record; state withheld at this stage due to regulatory concern). Australian specimens (NY91, NY92) represented adult and juvenile taken from the same laboratory colony maintained at Queensland University of Technology. Pakistan material (NY93) was collected from sugarcane in Multan, Punjab. All specimens were preserved in 95% ethanol and stored at −20 °C until processing.

### 2.2 Morphological character evaluation

All specimens were slide-mounted using the methodology described by Ahmed et al. (2025). Morphological characters were examined from slide-mounted adult females following Brookes (1978), Williams (2004), Schutze et al. (2019), Hernandez Europa et al. (2026), and Powell and Hauxwell (2026). Specimens from Australia, Barbados, Pakistan, and the U.S. consistently separated into two forms: the historical Type A lineage, which retains conspicuous translucent pores on the hind tibiae, and the invasive Type B lineage, in which these pores are absent. Type B specimens also match the new variant described by Schutze et al (2019), Hernandez-Europa et al (2026), and Powell and Hauxwell (2026), characterized by only two conical setae in the anal lobe cerarius, two size classes of quinquelocular pores, and poorly developed or absent tibial translucent pores, among other differences. Diagnostic traits evaluated included 1) the presence or absence of obvious translucent pores on the hind tibiae, 2) the number and arrangement of conical setae in the anal lobe cerarius, 3) size classes and distribution of quinquelocular pores on venter and dorsum, 4) density and distribution of crateriform oral rim tubular ducts, 5) presence, size, and distribution of slender spermatozoid ducts associated with crateriform ducts, 6) structure of lanceolate and flagellate setae across abdominal segments, 7) shape and development of the circulus and intersegmental line, 8) antennal segmentation and proportional lengths of segments I to IX, 9) claw morphology including presence of a denticle, and 10) arrangement of dorsal and ventral oral collar tubular ducts. These characters were compared across specimens from Barbados, Australia, Pakistan, and the U.S.

### 2.3 DNA extraction, Polymerase Chain Reaction (PCR), and sequencing

Genomic DNA was extracted individually using the DNeasy Blood and Tissue Kit (Qiagen, Maryland, U.S.) following the manufacturer’s protocol. DNA concentration and purity were assessed using a NanoDrop One spectrophotometer (Thermo Fisher Scientific, U.S.). Extracts were stored at −20 °C.

A partial fragment of the COI gene was amplified using the universal primers C1-J-2183 (Jerry; 5′-CAACATTTATTTTGATTTTTTGG-3′) and TL2-N-3014 (Pat; 5′-TCCAATGCACTAATCTGCCATATTA-3′), which target a mid-COI fragment overlapping the 3′ portion of the gene (Simon et al. 1994). Although the Folmer COI-5P barcode region (Hebert et al. 2003) is widely used in many insect groups, Folmer primers have very low amplification success in mealybugs and have consistently failed in our previous studies (Ahmed et al. 2015a,b). For this reason, and because the large mealybug taxa represented in GenBank are predominantly sequenced using Jerry–Pat, we selected this fragment as the most reliable and convenient COI region for Pseudococcidae. This ensured consistent amplification and direct comparability with existing reference datasets.

Each 25 µL reaction contained 12.5 µL Terra PCR Direct Buffer 2× (Takara Bio, California, U.S.), 0.5 µL Terra Polymerase Mix, 0.8 µL of each primer at 10 µM, 4 µL genomic DNA (20–50 ng/µL), and 6.4 µL nuclease-free water. The master mix composition followed the preparation procedures described in Ahmed et al. (2026). PCR amplification followed Ahmed et al. (2015a,b) and consisted of an initial denaturation at 95 °C for 3 min, followed by 40 cycles of 94 °C for 1 min, 57 °C for 30 s, and 72 °C for 1 min, with a final extension at 72 °C for 5 min.

Amplicons were visualized on 1.5% agarose gels stained with ethidium bromide. Products of the expected size were purified using NucleoSpin Gel and PCR Clean-up Kit (Macherey-Nagel, Pennsylvania, U.S.) and sequenced bidirectionally at Eurofins Genomics (Kentucky, U.S.). Raw chromatograms were inspected manually to ensure high-quality base calls across both strands.

### 2.4 NUMT screening

To verify that all newly generated COI sequences represented functional mitochondrial DNA rather than nuclear mitochondrial pseudogenes (NUMTs), each sequence was screened using a reproducible workflow implemented in R version 4.6.1 (R Core Team, 2026) in RStudio 2026.05.0+218 (“Golden Wattle”; Posit Software, Boston, U.S.). We conducted analyses using the *Biostrings* package within the Bioconductor framework (Huber et al. 2015).

We imported sequences as DNAString objects and cleaned them by removing gaps and ambiguous characters (N, ?, –). We translated the cleaned sequences using the invertebrate mitochondrial genetic code (NCBI translation table 5), the correct translation scheme for Hemiptera and other arthropods. This genetic code differs from the universal nuclear code in several codon assignments (e.g., TGA encodes tryptophan rather than a stop codon), making it essential for accurate mitochondrial COI translation in Sternorrhyncha.

Translated amino-acid sequences were examined for internal stop codons, frameshifts, and ambiguous residues. Codons containing unresolved or mixed bases were translated as “X”, indicating an ambiguous amino acid. Stop codons or X residues indicate potential NUMT origin or sequencing artifacts. The complete R script used for NUMT screening is provided in Supplementary File S1.

Following NUMT verification and sequence cleaning (Supplementary File S2), all newly generated COI sequences were trimmed to remove low-quality terminal bases and to reconstruct a single homologous region suitable for downstream analyses. This trimming step produced a consistent 708 bp fragment across all samples, ensuring positional homology and preventing inclusion of partial or misaligned sequence ends (Supplementary File S3). All newly generated COI sequences have been deposited in GenBank (accessions: XXXX–XXXX).

### 2.5 Reconstruction of homologous COI sequence regions

After NUMT verification and trimming, all sequences were then aligned against the complete mitochondrial genome of *Pl. citri* (NCBI accession PZ166947.1, UNVERIFIED: *Pl. citri* mitochondrion, complete genome) using MAFFT v7 (Katoh & Standley 2013) under default parameters (Supplementary File S4). Aligning to a full mitogenome reference allowed precise localization of the amplified fragment within the COI gene and ensured that all sequences corresponded to the same mitochondrial coordinates. This procedure reconstructed a single homologous COI region across all samples and prevented inclusion of partial, truncated, or mispositioned fragments. The final reference-anchored region was 708 bp for all sequences.

### 2.6 Verification of positional homology

To confirm that trimming and alignment preserved codon structure, we re-translated the final 708 bp reference-anchored COI region under the invertebrate mitochondrial genetic code (NCBI translation table 5) using the *Biostrings* package in Bioconductor (Huber et al. 2015). Translation of the reconstructed region verified correct codon phasing and the absence of frameshifts, internal stop codons, or ambiguous residues. These results demonstrate that positional homology was maintained throughout trimming and alignment and that the final COI region remains consistent with the NUMT-free status identified during initial screening (Supplementary File S2).

### 2.7 Sequence processing, BLAST retrieval, trimming, redundancy screening, and outgroup selection

All 15 newly generated COI sequences, after NUMT screening (Section 2.4), trimming and mitogenome anchoring (Section 2.5), and verification of positional homology (Section 2.6), were used as BLAST queries to retrieve homologous reference sequences from NCBI GenBank (Supplementary File S5). A total of 165 COI sequences were analyzed, including the 15 newly generated sequences and 150 GenBank records. Of the GenBank dataset, 142 sequences represented ingroup Pseudococcidae taxa and eight sequences represented non-Pseudococcidae outgroups selected according to the family-level phylogenomic framework of Deng et al. (2025), which provides the most robust evolutionary placement of Coccoidea families to date. Because initial BLAST hits ranged from ∼780–800 bp, only sequences with 100% query coverage, full positional homology to the reconstructed 708 bp region, and correct taxonomic assignment were retained (Pentinsaari et al. 2020). This filtering ensured that all external sequences corresponded precisely to the COI region amplified in this study and prevented inclusion of partial, truncated, or misannotated records. Metadata for all newly generated samples are provided in Table S1.

In GenBank, COI records for *Heliococcus* are limited and consist almost entirely of short fragments ranging from 340 to 360 bp. These include *H. singularis* (MG887770; 354 bp), *H. kurilensis* (MW881809; 354 bp), *H. summervillei* (MG833847 and MG833848; 355 bp), *H. buteae* (MG182699; 350 bp), and *H. bohemicus* (GU134695 and HM156737; 342–350 bp).

Because these fragments are less than half the length of the 708 bp homologous COI region reconstructed in this study, they could not be aligned to the full Jerry/Pat region, could not be translated reliably for NUMT screening, and produced non-overlapping alignments that inflated genetic distances. For these reasons, all short *Heliococcus* fragments were excluded. The *Heliococcus* sequences used in this study therefore consist of newly generated full-length Jerry/Pat sequences (NY73–NY76, NY84–NY93, NY99) and the single full-length GenBank record available for *Heliococcus* (Table S2).

To eliminate redundancy and ensure balanced representation across taxa, all sequences were screened for 100% identity using a custom R workflow implemented with the *ape* package (Paradis & Schliep 2019) using R version 4.6.1 (R Core Team, 2026) in RStudio version 2026.05.0+218 (“Golden Wattle”; Posit Software, Boston, U.S.). Pairwise raw distances were computed using dist.dna (model = “raw”), where a distance of 0 indicates complete identity across the aligned region. The script iteratively grouped sequences into identity clusters by selecting each unassigned sequence and retrieving all sequences with zero pairwise divergence. The complete R script used to screen for identical sequences and generate the 66 unique COI haplotypes is provided in Supplementary File S6.

The initial COI fragment was approximately 800 bp. To ensure positional homology across all samples and remove terminal regions with inconsistent read quality, sequences were uniformly trimmed to 708 bp as described in Section 2.5. All sequences were aligned using MUSCLE (Edgar 2004) implemented in MEGA 11 (Tamura et al. 2021). Translation-based verification of mitochondrial origin and positional homology, performed first during NUMT screening in Section 2.4 and again after trimming and alignment in Section 2.6, confirmed that the reconstructed region represented functional mitochondrial COI. The resulting 66 unique COI haplotypes (708 bp) generated by this workflow are provided in Supplementary File S7.

This procedure produced 58 non-redundant identity-based haplotype groups from the 157 ingroup Pseudococcidae sequences, preserving all unique sequence variation while preventing overweighting of identical sequences in downstream phylogenetic, genetic network, and genetic distance analyses (Table S2).

The GenBank dataset included representatives of multiple mealybug genera (*Balanococcus*, *Cataenococcus*, *Crisicoccus*, *Dysmicoccus*, *Ferrisia*, *Palmicultor*, *Paracoccus*, *Phenacoccus*, *Planococcus*, *Pseudococcus*, *Saccharicoccus*, *Tridiscus*) as well as eight non-Pseudococcidae scale insects used as outgroups. To avoid ambiguity among genera beginning with *P*, the following standardized abbreviations are used throughout: Pl. = *Planococcus*, Ph. = *Phenacoccus*, Pa. = *Paracoccus*, Ps. = *Pseudococcus*, and Pm. = *Palmicultor*. Outgroup selection followed the family-level phylogenomic framework of Deng et al. (2025), which provides the most comprehensive and robust evolutionary placement of Coccoidea families to date. The eight outgroups included *Acanthococcus lagerstroemiae* (Kuwana) (MZ312637), *Coccus hesperidum* Linnaeus (PP855271), *Dactylopius opuntiae* (Cockerell) (PP946188), *Fiorinia phantasma* Cockerell & Robinson (PQ846839), *Icerya purchasi* Maskell (PV738887), *Matsucoccus alabamae* Morrison (MT621224), *Orthezia urticae* (Linnaeus) (OL343331), and *Rhizoecus hibisci* (now placed in *Ripersiella hibisci* (Kawai & Takagi) (OQ833548). Incorporating these taxa ensured accurate rooting of the phylogeny and allowed lineage-level divergence within *H. summervillei* to be interpreted within a well-supported evolutionary context. Although COI is not appropriate for resolving deep family-level relationships, our analyses did not use COI for higher-level phylogeny. Instead, COI was applied only to root the ingroup topology and to verify that our *H. summervillei* haplotypes fall within the expected Pseudococcidae ingroup (Hebert et al. 2003; Armstrong & Ball 2005; Malausa et al. 2011; Downie & Gullan 2004; Ren et al. 2018).

Public COI records were reviewed for possible misidentifications following established practices in molecular diagnostics (Song et al. 2008; Foottit et al. 2008; Boykin et al. 2012; Ahmed et al. 2015a, 2015b, 2023, 2026). Sequences showing discordant placement in the COI phylogeny or unusually high divergence relative to their nominal species were flagged as potential misassignments or NUMTs and interpreted cautiously. This criterion has been used previously to identify misassigned GenBank accessions and NUMTs in Hemiptera, and we applied it consistently across analyses.

### 2.8 Maximum Likelihood phylogeny

Phylogenetic relationships among COI haplotypes were inferred using Maximum Likelihood (ML) in MEGA 11 (Tamura et al. 2021). Analyses were conducted on the 708 bp trimmed alignment using all 66 unique haplotypes derived from the full dataset. Model testing identified GTR+G as the best-fit nucleotide substitution model based on the lowest AICc and BIC values (Table S3). Maximum Likelihood trees were reconstructed under the GTR+G model using two discrete Gamma rate categories, and node support was evaluated using 1,000 bootstrap replicates.

Outgroup rooting followed Deng et al. (2025). These outgroups were included solely to stabilize tree rooting and to confirm that all *H. summervillei* haplotypes fall within the expected Pseudococcidae clade.

The purpose of the phylogenetic analysis was to evaluate species-level cohesion within *H. summervillei*, not to reconstruct broader Pseudococcidae relationships. Specifically, the ML tree was used to test whether (i) Type A and Type B haplotypes form a single, well-supported clade, (ii) Type A haplotypes group tightly within a single sub-clade, and (iii) both mitochondrial lineages occupy the correct phylogenetic position relative to the surrounding Pseudococcidae ingroup. The collapsed ML phylogeny is shown in Figure 1, and the complete haplotype-level ML tree is presented in Figure S1.

**FIGURE 1.**
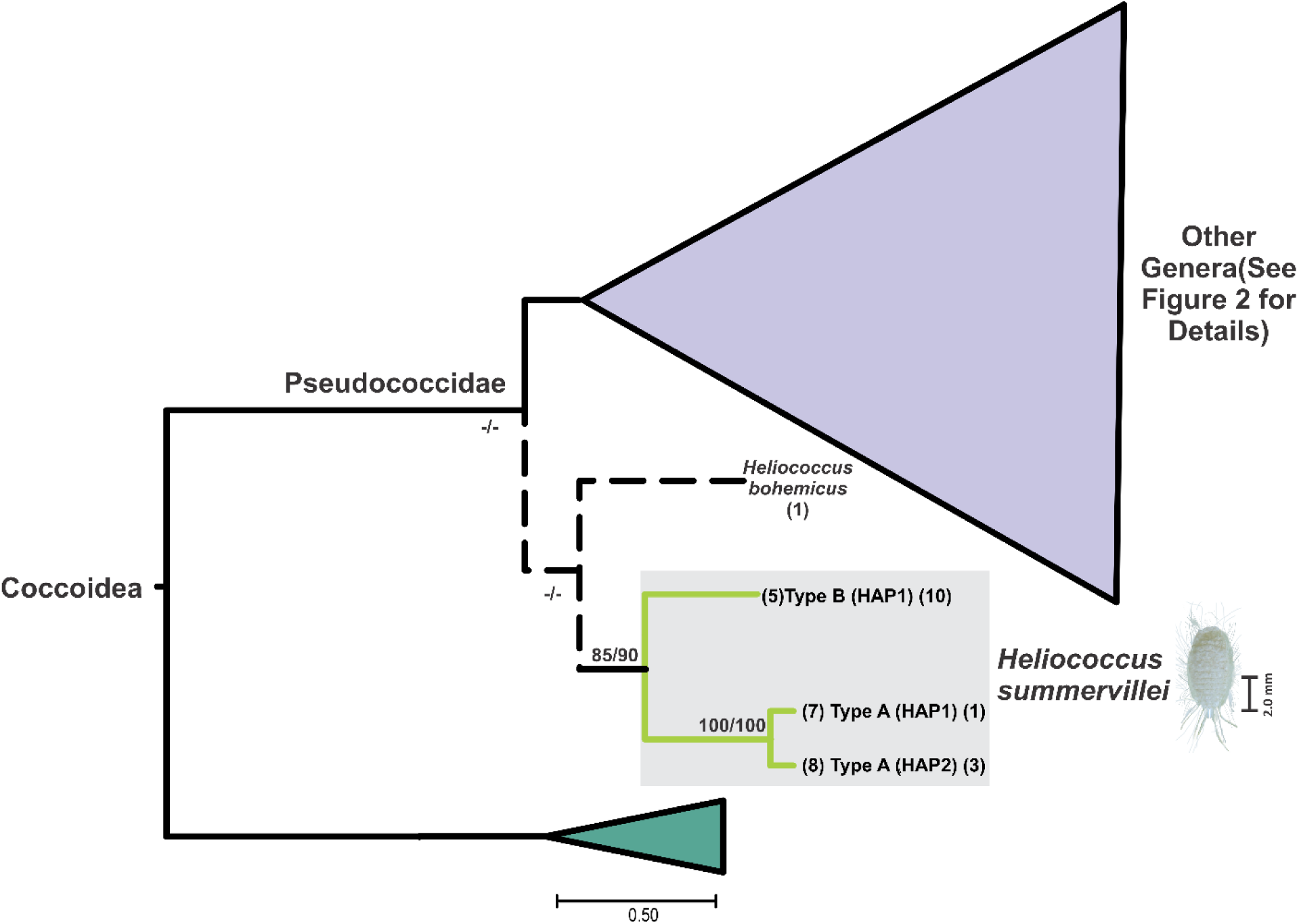
Maximum Likelihood (ML) phylogeny of *Heliococcus summervillei* COI (Cytochrome Oxidase I) haplotypes derived from the 708 bp COI alignment. Individual haplotypes were collapsed into identity-based groups (see details in Table S2) to improve visual clarity. The *H. summervillei* clade is well-supported in both analyses, with the combined Type A + Type B lineage receiving strong support (ML bootstrap = 85; IQ-TREE UFBoot = 100). Within this clade, the Type A haplotypes (Barbados; Groups 7–8) form a deeply divergent subclade that is fully supported in both frameworks (ML bootstrap = 100; SH-aLRT = 100; UFBoot = 100). The *H. bohemicus* clade is shown with dashed outlines because this taxon is placed outside its expected position in the full, uncollapsed phylogeny (Supplementary Figure S1). Dashed lines indicate taxa whose placement differs from their expected clade when branch lengths are fully represented. Branch lengths are drawn to scale, using the same scaling applied in the full ML tree (via the “toggle scaling” option). Because the tree is collapsed, branch lengths represent the underlying ML topology. The complete, uncollapsed haplotype-level phylogeny, including full branch lengths, all outgroup placements, and all misplacement indicators, is provided in Supplementary Figure S1. Bootstrap values < 50 were replaced with a dash (–) for clarity.

An additional ML phylogeny was generated using IQ-TREE v1.6.11 (Nguyen et al. 2015; Trifinopoulos et al. 2016; Minh et al. 2020) to provide a more robust assessment of branch support and to follow current best practices in molecular systematics. ModelFinder (Kalyaanamoorthy et al. 2017) identified TIM+F+R5 as the best-fit substitution model under BIC. Rate heterogeneity was modeled using the FreeRate approach with five rate categories. Node support was evaluated using 10,000 ultrafast bootstrap replicates (UFBoot2; Hoang et al. 2018) and 1,000 SH-aLRT single-branch tests (Guindon et al. 2010). The IQ-TREE analysis produced two trees: (i) a branch-length ML tree with SH-aLRT and UFBoot support values, and (ii) a bootstrap support tree with branch lengths scaled to genetic distances. Outgroup rooting used the same designated outgroup taxa (alignment positions 59–66). Both IQ-TREE phylogenies are provided in the supplementary materials (Figure S2), and all IQ-TREE model-selection and support statistics are reported in Supplementary File S8.

### 2.9 Genetic network analysis

To visualize genetic relationships among all COI haplotypes, including both ingroup Pseudococcidae and the eight non-Pseudococcidae outgroup taxa, a statistical parsimony network was constructed in TCS 1.23 (Clement et al. 2000) under the Templeton 95% connection limit using an aligned sequence FASTA file in Supplementary File S5. Analyses were performed on the 708 bp trimmed alignment using the 58 identity-based haplotype ingroups and 8 outgroup haplotypes defined in Table S2. Inclusion of outgroup haplotypes allowed the network to represent the full mitochondrial divergence structure across Coccoidea and provided a direct comparison between *H. summervillei* lineages and deeper family-level splits. This approach follows the genetic-network framework applied by De Barro & Ahmed (2011), who demonstrated that genetic networks can resolve cryptic species boundaries, identify lineage-level divergence, and reveal invasion-related mitochondrial structure within the *B. tabaci* species complex. Applying the same logic here enables assessment of whether divergence between the historical Type A and invasive Type B lineages of *H. summervillei* falls within typical intraspecific ranges or approaches thresholds associated with species-level differentiation. The resulting haplotype-level network, showing both ingroup clusters and outgroup placements, is presented in Figure 2.

**FIGURE 2.**
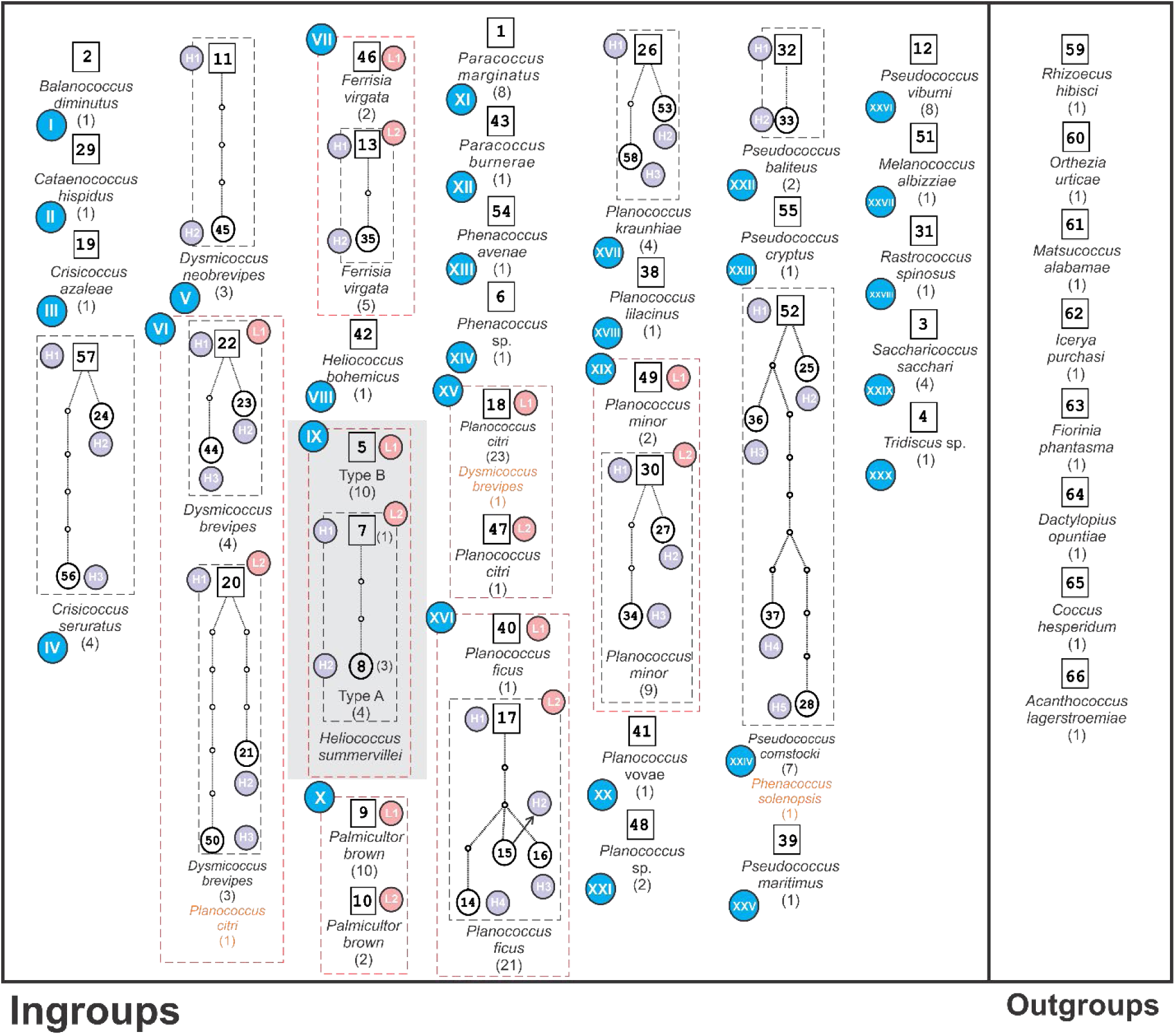
Statistical parsimony haplotype network (TCS v1.23, Templeton 95% connection limit) illustrating mutational relationships among *Heliococcus summervillei* COI (Cytochrome Oxidase I) haplotypes within the broader Pseudococcidae dataset. Blue circles represent species-level identity groups labeled with Roman numerals corresponding to Table S2. Purple circles denote *H. summervillei* haplotypes, and pink circles indicate lineage-level clusters within other species. Boxed numeric labels correspond to identity-based haplotype groups used in the collapsed phylogeny (Figure 1; the full, uncollapsed species-level phylogeny corresponding to these groups is provided in Supplementary Figure S1). Black dashed boxes outline mitochondrial lineages, and red dashed boxes indicate species-level clusters. Species names shown in black represent taxa clustering together as expected, whereas species names shown in orange indicate taxa misidentified or mislabeled in GenBank. Numbers in parentheses denote the total sequences represented by each species or lineage. Ingroup Pseudococcidae and outgroup Coccoidea taxa are displayed separately to provide phylogenetic context.

### 2.10 Genetic distance analysis and Principal Coordinates Analysis

Pairwise Kimura 2-Parameter (K2P) genetic distances were calculated in MEGA 11 (Tamura et al. 2021) using the 708 bp trimmed COI- alignment (Table S4). Distances were computed among all 58 identity-based haplotype groups, enabling direct comparison of divergence levels across *H. summervillei* and related Pseudococcidae taxa. The complete set of K2P values is provided in Table S4 and served as the source dataset for all downstream multivariate analyses.

Outgroup COI sequences were excluded from K2P distance calculations because they did not overlap the 708 bp homologous region reconstructed for ingroup taxa. When forced into the alignment, these non-overlapping regions produced artificially inflated divergence estimates, including values exceeding 1.0. For this reason, all outgroup sequences (OQ833548, OL343331, MT621224, PV738887, PQ846839, PP946188, PP855271) were removed from genetic distance and PCoA analyses. Only *A. lagerstroemiae* (MZ312637), which retained sufficient overlap with the homologous region, was included in the K2P distance matrix. All outgroups, including *Acanthococcus*, were retained in phylogenetic and genetic network analyses because these methods do not require full positional homology across the 708 bp region, and partial overlap is sufficient for accurate rooting and for visualizing deep mitochondrial divergence.

To evaluate lineage-level separation within *H. summervillei*, K2P distances were specifically examined between haplotypes representing the historical Type A lineage (Barbados) and those corresponding to the invasive Type B lineage (Australia, Pakistan and U.S.). These values were compared with divergence observed among other mealybug genera and the non-Pseudococcidae outgroup taxa, providing a broader evolutionary context for assessing whether mitochondrial divergence within *H. summervillei* fell within typical intraspecific ranges or approached interspecific thresholds.

For multivariate visualization of genetic structure, the K2P distances exported from MEGA were reformatted into square distance matrices using the reproducible workflows in Supplementary Files S9 and S10. The full matrix, including all 58 ingroup haplotypes and the eight outgroup taxa, is provided in Table S4, whereas the ingroup-only matrix used for lineage-level ordinations is provided in Supplementary File S11. Two complementary ordinations were generated. First, a Principal Coordinates Analysis (PCoA) including all ingroup Pseudococcidae haplotypes and the outgroup taxa [*A. lagerstroemiae* (MZ312637)] was performed using the reproducible R workflow in Supplementary File S10 with R version 4.6.1 (R Core Team, 2026) in RStudio version 2026.05.0+218 (“Golden Wattle”; Posit Software, Boston, U.S.). This analysis (Figure 3A) illustrates the strong separation between Pseudococcidae ingroups, which form a tight cluster, and the non-Pseudococcidae outgroups, which fall far to the right along the major axis of variation.

**FIGURE 3.**
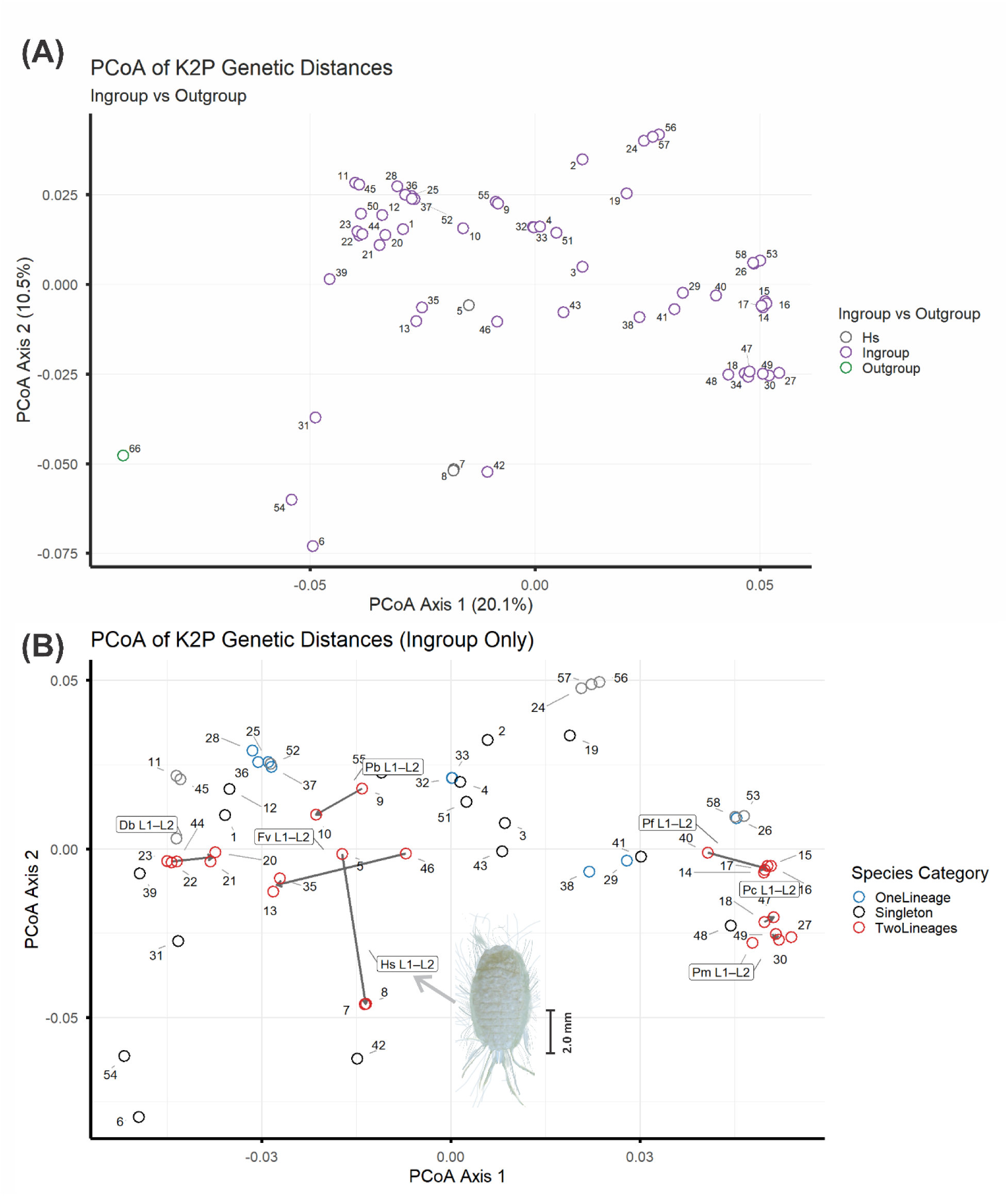
Principal Coordinates Analysis (PCoA) of K2P genetic distances for COI (Cytochrome Oxidase I) haplotypes. (A) PCoA of 59 haplotypes, including 58 ingroup Pseudococcidae and one non-Pseudococcidae outgroup (Hap66). Hollow circles represent individual haplotypes, colored by group: ingroup (purple), outgroup (green), and *Heliococcus summervillei* (Hs; gray). Ingroup haplotypes form a tight cluster with shallow genetic distances (0.07–0.15 K2P), whereas outgroups exhibit deep divergence (∼1.0–1.5 K2P) and separate strongly along Axis 1. This ordination provides a broad view of mitochondrial divergence across Pseudococcidae and highlights the clear separation between ingroup taxa and the selected outgroup. (B) Principal Coordinates Analysis (PCoA) of K2P genetic distances for all ingroup COI haplotypes. Hollow circles represent individual haplotypes, colored by species category: singletons (black), species with one mitochondrial lineage (blue), and species with two lineages (red). Arrows connect lineage centroids within species to illustrate the magnitude of mitochondrial divergence. Lineage comparisons are labeled using genus–species initials followed by lineage identifiers: Fv (*Ferrisia virgata*), Pb (*Palmicultor browni*), Pc (*Planococcus citri*), Pf (*Planococcus ficus*), Pm (*Planococcus minor*), Hs (*Heliococcus summervillei*), and Db (*Dysmicoccus brevipes*). Species with shallow intraspecific divergence (<3%) form tight clusters, whereas species with two lineages show deep mitochondrial splits (0.15–10%). *H. summervillei* Type A and Type B (Hs L1–L2) exhibit ∼10% divergence and appear as distinct clusters. NY77 (*Phenacoccus* sp.) is a distant outlier (∼18.45%) and falls outside the *H. summervillei* cluster.

Second, an ingroup-only PCoA was generated using the workflow in Supplementary File S10, which imports the ingroup-only K2P matrix from Supplementary File S11 and resolves lineage-level structure within Pseudococcidae (Figure 3B). These scripts imported the finalized K2P matrices, applied centering and scaling, and performed eigenvalue decomposition to extract orthogonal axes summarizing the major components of genetic variation.

All ordinations were implemented using the *ape* package (Paradis & Schliep 2019) for PCoA computation, *ggplot2* (Wickham 2016) for graphical rendering, *ggrepel* (Slowikowski et al. 2026) for non-overlapping labels, and *dplyr* (Wickham et al. 2023) for data handling and category assignment. The resulting PCoA ordinations were used to quantify clustering among haplotypes and evaluate whether patterns of mitochondrial variation aligned with established lineage assignments (Type A, Type B, excluded lineage).

PCoA was performed following standard procedures (Gower 1966; Legendre & Legendre 2012; Anderson & Willis 2003). The distance matrix was centered prior to eigenvalue decomposition, and negative eigenvalues were inspected and found to have negligible influence on the ordination. Variance explained by each axis was calculated from the eigenvalues, and graphical adjustments were made only to improve clarity without altering the underlying analysis (Borcard et al. 2018).

## 3 Results

### 3.1 Morphological differentiation of lineages

Morphological examination demonstrated that the Barbados specimens correspond to the holotype described by Brookes (1978) (also known as Type A). These individuals exhibited translucent pores on the hind tibiae, three conical setae in the anal-lobe cerarius, a single size class of quinquelocular pores, fully developed crateriform oral-rim tubular ducts with associated spermatozoid ducts, lanceolate dorsal setae, flagellate ventral setae, a claw with a denticle, and a circulus with a distinct intersegmental line. These characters match the original species description and subsequent accounts from early outbreak regions.

In contrast, all invasive-range specimens matched the Type B variant documented in recent Australian and U.S. outbreaks. These individuals lacked or had very poorly developed tibial translucent pores, possessed two conical cerarius setae, exhibited two size classes of quinquelocular pores, showed reduced crateriform duct density, and displayed modified spermatozoid ducts and altered antennal proportions consistent with Schutze et al. (2019), Hernandez Europa et al. (2026), and Powell & Hauxwell (2026). Despite these differences, both variants retained most of the diagnostic suite of characters defining *H. summervillei* (Hernandez Europa et al. 2026). These morphological patterns suggest that the two mitochondrial lineages correspond to historically stable morphological variants, although their species status remains unresolved.

One Barbados specimen (NY77) did not match either Type A or Type B. Instead, its morphology aligned with *Phenacoccus* based on the diagnostic characters in Ahmed et al. (2025), including the absence of ventral quinquelocular pores, the presence of dorsal multilocular pores arranged in rows across abdominal segments, the lack of auxiliary setae in cerarii anterior to the anal-lobe cerarius, and a cerarius count exceeding ten pairs. NY77 also lacked all defining traits of *Heliococcus*, such as crateriform tubular ducts, spermatozoid ducts, tibial translucent pores, lanceolate dorsal setae, and the distinctive circulus with an intersegmental line (Schutze et al. 2019; Hernandez Europa et al. 2026; Powell & Hauxwell 2026). These characters are not merely inconsistent with *Heliococcus*, they are diagnostically incompatible with the genus, demonstrating that NY77 is a *Phenacoccus* species.

### 3.2 Sequence recovery, trimming, and identity-based haplotype grouping

All 15 newly generated COI sequences were successfully recovered and translated without internal stop codons, confirming their mitochondrial origin. BLAST comparisons identified close matches across multiple Pseudococcidae genera, and 150 taxonomically verified GenBank sequences were incorporated into the final dataset (Table S1–2).

Redundancy screening reduced the full dataset to 66 unique COI haplotypes (Supplementary File S7), representing all distinct mitochondrial variants present across ingroup and outgroup taxa. These haplotypes captured the complete sequence diversity available for downstream phylogenetic, genetic network, and genetic-distance analyses.

Identity-grouping of all sequences yielded 66 non-redundant identity-based haplotype groups (Table S2). These groups included representatives of both *H. summervillei* lineages, multiple Pseudococcidae genera, and eight non-Pseudococcidae outgroup taxa, providing a phylogenetically balanced framework for evaluating lineage-level divergence.

Within *H. summervillei*, three identity groups contained all focal specimens (Table S1–2). One group corresponded to the invasive lineage, whereas two groups represented the historical lineage. A fourth group contained a deeply divergent specimen (NY77). This structure revealed two stable mitochondrial haplotypes within the Barbados Type A lineage and one haplotype representing the invasive Type B lineage, along with a single outlier (NY77) indicative of a distinct species-level lineage.

### 3.3 Maximum Likelihood phylogeny

Maximum Likelihood analysis of the 66 unique COI haplotypes recovered a well-resolved topology that clearly separated ingroup Pseudococcidae from the eight non-Pseudococcidae outgroups (Figure 1; Figure S1). The best-fit substitution model (GTR+G) produced a stable tree structure with consistent placement of major genera and support for the *H. summervillei* clade. Within this clade, both mitochondrial lineages were recovered together (bootstrap = 85), and the two Barbados Type A haplotypes formed a fully supported subclade (bootstrap = 100). The invasive Type B haplotype remained uniform and clustered immediately adjacent to Type A, reflecting their close mitochondrial affinity.

The ML topology also revealed two anomalous placements. Two *Pseudococcus* haplotypes were positioned inside the *Dysmicoccus brevipes* (Cockerell) clade, and *H. bohemicus* appeared within the *Phenacoccus* cluster. These anomalies were limited to a small number of haplotypes and did not affect the overall recovery of major familial boundaries.

Branch-length reconstructions (Figure S1) produced the same lineage structure and further highlighted the deep divergence between *H. summervillei* and other genera. Long internal branches separated *H. summervillei* from neighboring taxa, and the split between Type A and Type B was consistently recovered in the ML topology. Together, these results confirm the distinct mitochondrial lineages within *H. summervillei* and their stable placement within Pseudococcidae.

The IQ-TREE ML phylogeny produced a stable topology that clearly separated most Pseudococcidae ingroup taxa from the designated outgroups, although two outgroup sequences (61 and 66) were recovered within the ingroup with weak support (Figure S2). Within *H. summervillei*, both mitochondrial lineages again clustered together, with the two Type A haplotypes forming a fully supported subclade (100) and the combined Type A + Type B lineage receiving support (90). As in the MEGA ML tree, a small number of haplotypes were placed within an unexpected cluster, but these placements were weakly supported and did not affect higher-level relationships. *Heliococcus bohemicus* was positioned inside the *Phenacoccus* group with support of 94, although its overall placement in the tree was not well supported (Figure S2A). Branch-length patterns further highlighted the deep divergence between *H. summervillei* and neighboring genera, and both the ML and consensus bootstrap trees consistently recovered the split between Type A and Type B (Figure S2B). Overall, the IQ-TREE results corroborate the MEGA ML findings and confirm that the newly generated haplotypes fall within Pseudococcidae and form two closely related mitochondrial lineages (Figure S1 and S2).

### 3.4 Statistical parsimony genetic network and genetic distances analysis

The statistical parsimony network was constructed using the 66 identity-based COI haplotypes defined in Table S2. Seventeen sequences were represented by a single haplotype and showed no intraspecific variation in genetic network analysis: identity groups I (2), II (29), III (19), VIII (42), XI (1), XII (43), XIII (54), XIV (6), XVIII (38), XX (41), XXI (48), XXIII (55), XXV (39), XXVI (12), XXVII (51), XXVIII (31), and XXX (4) (Figure 2). We excluded these singletons from lineage and haplotype comparisons because they cannot inform intralineage divergence (Figure 2). Singleton haplotypes refer to identity groups represented by a single unique COI haplotype, regardless of the number of sequences contained within that group.

Multiple species exhibited one mitochondrial lineage with several haplotypes and uniformly shallow intraspecific distances. *Dysmicoccus neobrevipes* Beardsley (V) (*n* = 3) contained two haplotypes, *Pseudococcus baliteus* Lit (XXII) (*n* = 2) contained two haplotypes, *Crisicoccus seruratus* (Kanda) (IV) (*n* = 4) contained three haplotypes, *Planococcus kraunhiae* (Kuwana) (XVII) (*n* = 4) contained three haplotypes, and *Pseudococcus comstocki* (Kuwana) plus *Ph. solenopsis* (XXIV) contained five haplotypes (*n* = 5). Within XXIV, haplotype 28 (H5) was misidentified in GenBank as *Ph. solenopsis*, while the remaining haplotypes matched *P. comstocki*. Pairwise distances among haplotypes in these one-lineage species ranged from 0.001– 0.03, with *Ps. comstocki* specifically showing 0.001–0.014 (∼1.4%) (Figure 2).

Seven species contained two mitochondrial lineages. *Ferrisia virgata* (Cockerell) (VII) had L1 (46) (L1 contained 2 *F. virgata* sequences, *n* = 2) and L2 (13, 35) (*n* = 5); intralineage distances within L2 were 0.001 (0.14%), and interlineage distances were 0.085–0.086 (8.5– 8.6%). *Palmicultor browni* (Williams) (X) had L1 (9) (*n* = 10) and L2 (10) (*n* = 2), separated by 0.046 (4.6%). *Planococcus. citri* (XV) had L1 (18) (*n* = 24, including one misidentified *D. brevipes* sequence) and L2 (47) (*n* = 1); the distance between 18 and 47 was 0.0015 (∼0.15%). *Planococcus ficus* (Signoret) (XVI) contained one lineage represented by haplotype 40 (*n* = 1) and a second lineage represented by four haplotypes (14–17) (*n* = 21); intralineage distances within the multihaplotype lineage were 0.003–0.006 (0.2–0.6%), and interlineage distances were 0.037–0.040 (∼3.7–4.0%). Because one lineage was represented by a single haplotype, all interlineage comparisons involved that single sequence, while the multihaplotype lineage contained several distinct haplotypes. *Planococcus minor* (Maskell) (XIX) contained L1 (49) (*n* = 2) and L2 (30, 27, 34) (*n* = 9); intralineage distances within L2 were 0.0014–0.0057 (∼0.14– 0.57%), and interlineage distances were 0.0029–0.0044 (∼0.3–0.4%). Because L1 was represented by a single haplotype, all interlineage comparisons involved that single sequence, whereas L2 contained multiple distinct haplotypes, resulting in a broader intralineage range despite overall lower divergence. *Dysmicoccus brevipes* (VI) contained two mitochondrial lineages. L1 included haplotypes 22, 23, and 44 (distances 0.0014–0.0288) (*n* = 4). L2 included haplotypes 20, 21, and 50 (distances 0.0014–0.0288) (*n* = 4), including one misidentification; haplotype 50 was misidentified as *Pl. citri*. Interlineage distances between L1 and L2 were 0.026–0.030 (0.026–0.030) (∼2.6–3.0%) (Figure 2).

*Heliococcus summervillei* (IX) contained two mitochondrial lineages. Lineage 1 (Type B) was represented by haplotype 5 (*n* = 10). Lineage A (Type A) was represented by haplotypes 7 and 8 (*n* = 4). Intralineage distances within Type A were 0.003 (0.28%). Interlineage distances between Type A and Type B were 0.0997–0.0999 (9.97–9.99%). No intralineage variation was observed within Type B. Misidentified sequences were excluded from lineage interpretation, including haplotype 28 (XXIV) (*Ps. comstocki*), haplotype 18 (XV) (*Pl. citri*), and haplotype 50 (VI) (*D. brevipes*) (Figure 2).

Across all ingroup species, intralineage distances were typically <0.03 (<3%), interlineage distances within species ranged from ∼0.0015–0.10 (0.15–10%), and interspecific distances were generally >0.11 (>11%).

### 3.5 Principal Coordinates Analysis and genetic distance analysis

Genetic distances (K2P) ranged from 0.000 to ∼0.12 for ingroup Pseudococcidae, whereas the designated outgroup exhibited much deeper divergence (0.16–0.21) (Table S4) Typical intraspecific values were <0.03 (<3%), and interspecific values were >0.11 (>11%). To visualize broad-scale divergence across all taxa, a PCoA including all ingroup mealybug haplotypes and the non-Pseudococcidae outgroup [*A. lagerstroemiae* (MZ312637)] was generated (Figure 3A). This ordination revealed a clear separation between Pseudococcidae and the non-Pseudococcidae outgroup, with all mealybug haplotypes, including both *H. summervillei* lineages, forming a tight, cohesive cluster characterized by shallow K2P distances (0.00–0.12). In contrast, the designated outgroup (*A. lagerstroemiae*) showed K2P divergence of 0.16–0.21 and occupied a distinct position relative to the ingroup cluster along PCoA Axis 1 (Figure 3A). The placement of *H. summervillei* within the mealybug cluster confirms its mitochondrial affinity with other Pseudococcidae.

K2P distances revealed three distinct divergence patterns within the *H. summervillei* complex. First, within-lineage divergence was extremely low. The invasive lineage (Hs L1) showed no divergence (= 0.0000) across Australia, Pakistan, and the U.S. (Florida, Louisiana, Texas, and an anonymous state). The Barbados lineage (Hs L2) contained two closely related haplotypes with only 0.000–0.003 divergence, reflecting extremely shallow mitochondrial variation within the lineage. These values correspond to the tight clusters observed in the PCoA ordination (Figure 3B), where both lineages group closely along PCoA Axis 2, show no internal structure, and are separated primarily along PCoA Axis 1.

Second, divergence between the Barbados and invasive lineages averaged ∼10% (0.09971– 0.09986). In the PCoA, this split is expressed as a clear separation along Axis 1, with an arrow connecting the lineage centroids (Hs L1–L2) (Figure 3B). Third, one specimen (Haplotype 6) (NY77) showed ∼18.45% divergence from both lineages (0.18441–0.18467), well within the interspecific range for mealybugs (Figure 3B).

Beyond *H. summervillei*, the PCoA also reflected lineage structure in other mealybug taxa included for comparative context. Species with one mitochondrial lineage formed tight clusters (blue), whereas species with two lineages formed distinct clusters connected by arrows (red). Examples include *F. virgata* (Fv L1–L2), *Pa. browni* (Pb L1–L2), *Pl. citri* (Pc L1–L2), *Pl. ficus* (Pf L1–L2), *Pl. minor* (Pm L1–L2), and *D. brevipes* (Db L1–L2). These species exhibited interlineage distances of ∼0.0015 to 0.10 (0.15–10%), matching the deep mitochondrial structuring seen in *H. summervillei* (Figure 3B).

Overall, the PCoA ordination (Figure 3A–B) visually reinforces the genetic-distance patterns by showing extremely shallow intralineage divergence, a deep but sub-interspecific split between Type A and Type B, and a clearly distinct outlier representing a different species. This combined evidence supports historically stable mitochondrial lineages within *H. summervillei* and highlights the broader pattern of deep lineage divergence across multiple Pseudococcidae.

## 4 DISCUSSION

This study provides the first COI sequences generated for *H. summervillei*, addressing a long-standing gap in molecular diagnostics for this invasive mealybug. Accurate mealybug identification is notoriously difficult because of cryptic morphology, reduced and overlapping characters, and the need for highly technical slide-mounted preparations (Williams & Granara de Willink 1992; Williams 2004; Malausa et al. 2011; Ahmed & Deeter 2022). COI is widely used in diagnostics and pest management because of its high amplification success and extensive reference coverage (Hebert et al. 2003; Armstrong & Ball 2005; García Morales et al. 2016; Ren et al. 2018). Generating COI sequences for both Type A and Type B variants therefore provides a molecular foundation for resolving lineage structure and supporting identification in quarantine, biosecurity, and applied management programs implemented directly in pastures, turfgrass systems, sod production fields, and managed landscape environments.

COI genetic distance data reveal two deeply diverged mitochondrial lineages: the Type A lineage, which contains two closely related haplotypes, and the invasive Type B lineage, which contains a single globally distributed haplotype. These mitochondrial divergence patterns corroborate the morphological evidence reported in earlier work, which consistently recognizes Type A and Type B variants within *H. summervillei* (Schutze et al. 2019; Hernandez-Europa et al. 2026; Powell & Hauxwell 2026). The ∼10% sequence divergence between these lineages exceeds typical intraspecific COI values reported for most mealybugs, yet remains below interspecific thresholds and falls within a range where species limits cannot be confidently inferred (Downie & Gullan 2004; Malausa et al. 2011; Ren et al. 2018). Ren et al. (2018) reported that most mealybug species exhibit <3% intraspecific divergence, but several taxa show values between 7–9%, and some morphologically identical lineages reach ≥8.5%, indicating that deep mitochondrial structuring can occur without clear species-level separation. These patterns indicate historically stable mitochondrial divergence between the Type A and invasive Type B lineages, while leaving their taxonomic status unresolved without a full integrative revision (Malausa et al. 2011; Downie & Gullan 2004; De Barro & Ahmed 2011; Hardy et al. 2008).

Hernandez Europa et al. 2026 established the morphological and endosymbiont separation of Type A and Type B in Australia, but that study did not generate or analyze COI sequences. To date we analyzed the data; none of the mitochondrial information presented here, including the full-length Jerry Pat COI sequences, the reconstruction of a homologous 708 bp region, the phylogenetic, genetic network, genetic distance estimates, or the documentation of a single invasive Type B haplotype, had been reported previously. The present study therefore provides the first COI sequences for *Heliococcus summervillei* and the first comparative mitochondrial assessment of Type A and Type B across multiple regions. These results complement the morphological and endosymbiont patterns described by Hernandez Europa et al. 2026, and extend that work by adding a mitochondrial dataset that was not part of the earlier study.

Although no historical Australian Type A specimen was available for COI sequencing, the Barbados lineage is provisionally assigned to Type A based on morphology. Barbados specimens exhibit the diagnostic tibial translucent pores, anal-lobe cerarius structure, and the number and size classes of quinquelocular pores that have consistently characterized Type A in the original description and subsequent Australian work (Schutze et al. 2019; Hernandez Europa et al. 2026; Powell & Hauxwell 2026). These traits provide the best available evidence for aligning the Barbados lineage with Type A. However, we acknowledge that this interpretation is based solely on morphology and may require revision as additional Australian Type A material becomes available for sequencing.

Across the broader dataset, deep mitochondrial structuring was common in our study. Several mealybug species contained multiple COI lineages with very low intralineage variation and moderate to high interlineage divergence, including *F. virgata*, *Pa. brown*, *Pl. citri*, *Pl. minor*, and *D. brevipes* (Figure 1–3). These patterns are consistent with previous reports of elevated mitochondrial divergence within morphologically defined mealybug species (Downie & Gullan 2004; Malausa et al. 2011; Ren et al. 2018). In our dataset, interlineage distances ranged from approximately 0.15–10%, while intralineage distances were uniformly low (<1–3%). Our PCoAs (Figure 3) further supported these patterns by clearly separating major COI lineages within each species and demonstrating that the genetic distance between the historical Type A and invasive Type B lineages of *H. summervillei* is comparable to, and in some cases greater than, lineage-level divergence observed in other Pseudococcidae. Such mitochondrial divergences within species are well documented across Hemiptera and Pseudococcidae, arising from processes such as incomplete lineage sorting, introgression, demographic isolation, and endosymbiont-associated sweeps (Hurst & Jiggins 2005; Dinsdale et al. 2010; Després 2019; Poveda-Martínez et al. 2020). Consequently, substantial COI divergence can occur within nominal species, and mitochondrial patterns alone should not be interpreted as species-level separation without considering them alongside biological data.

The ML phylogeny and parsimony haplotype network support this mitochondrial lineage divergence. Although a few haplotypes were placed outside their expected clades or networks, likely reflecting misidentifications in GenBank or the limited resolving power of COI, the major familial boundaries were recovered, and both *H. summervillei* lineages consistently formed a single clade within Pseudococcidae. Type A and Type B clustered together with strong bootstrap support. The two Type A haplotypes formed a fully supported subclade, whereas Type B remained uniform, reflecting the shallow intralineage variation documented in the distance analyses. Because COI represents a single mitochondrial marker, these phylogenetic patterns cannot resolve species boundaries; however, they demonstrate that both mitochondrial lineages belong to the same species-level clade and share a common evolutionary origin.

Evidence from endosymbionts provides a second, independent means of differentiating Type A and Type B. In a separate study, Hernandez-Europa et al. (2026) showed that Type A and Type B carry distinct *Tremblaya phenacola* (Buchner) Amplicon Sequence Variants (ASVs), with no overlap between lineages, a pattern that mirrors their consistent morphological separation. However, the same study found that ribosomal markers (18S, 28S-D2) showed limited and inconsistent differentiation between the two types, and the dynamin dataset included only a single usable Type A sequence, preventing meaningful comparison across lineages. These results suggest that some genomic differences may exist between Type A and Type B, but the available nuclear data remain incomplete and do not yet allow strong conclusions about the extent of their separation.

Although our morphological comparisons included original species descriptions, evaluation of slide-mounted specimens, and our molecular data show consistent mitochondrial divergence, this study does not attempt to revise the taxonomy of *H. summervillei*. A full taxonomic treatment incorporating detailed morphology, multilocus nuclear sequencing, endosymbiont characterization, and expanded geographic sampling will be required to determine whether the Type A and Type B lineages represent distinct species; however, this was outside the scope of this study.

The genetic structure documented in *H. summervillei* has practical implications. Genetic variation in mealybugs can influence insecticide susceptibility, host range, developmental biology, dispersal potential, and biological control compatibility (Downie & Gullan 2004; De Barro & Ahmed 2011; Malausa et al. 2011; Waqas et al. 2019; Shankarganesh et al. 2022; Kumar et al. 2023). Accurate matching of parasitoids to the correct host lineage is essential for effective classical biological control (Muniappan et al. 2006). Although functional differences among *H. summervillei* lineages were not evaluated here, recognizing distinct mitochondrial variants provides a basis for future work on lineage-specific management responses and may assist in reconstructing introduction pathways, which is critical for preventing secondary spread and informing regulatory and chemical-control decision-making frameworks.

The invasion history of Type B is consistent with a single, widely distributed invasive lineage. This same mitochondrial lineage was recovered in our study from Australia, Pakistan, and multiple U.S. states, demonstrating its broad geographic distribution. Its broad distribution and no intralineage COI variation suggest a single expanding invasive haplotype rather than multiple introductions of genetically distinct lineages. Similar single-lineage incursions have been reported in other invasive insects, including *Ph. solenopsis* during its expansion across Asia (Ahmed et al. 2015a), *T. parvispinus* during its spread across multiple continents (Ahmed et al. 2025b), and *Amrasca biguttula* (Ishida) following its introduction into the U.S. (Ahmed et al. 2026). Comparable lineage-specific patterns have also been documented in other invasive phytophagous insects, where population growth, host shifts, and insecticide exposure can shape genetic patterns relevant to management (De Barro & Ahmed 2011; Ahmed et al. 2015a; Xu et al. 2024; Ahmed et al. 2025b, 2026).

One specimen (NY77) fell far outside the *H. summervillei* clade and did not match *H. summervillei* morphologically. Its overall morphology was consistent with *Phenacoccus* rather than *Heliococcus*, and both phylogenetic reconstructions placed NY77 in a completely separate clade from the two *H. summervillei* mitochondrial lineages. In the genetic analyses, including the PCoA, NY77 formed a distant outlier far from both Hs L1 and Hs L2, reflecting its deep COI divergence. Although collected from lawn grasses alongside the Barbados lineage, its morphology and genetic distance indicate that NY77 represents a different species-level lineage. This case highlights how overlapping field habitats and cryptic external field traits can lead to misidentifications in mealybugs, and underscores the importance of COI confirmation in resolving species boundaries in groups like mealybugs (Malausa et al. 2011; Ahmed & Deeter 2022; Ahmed et al. 2025a).

Taken together, the evidence indicates two deeply diverged mitochondrial lineages associated with Type A and Type B, each supported by endosymbiotic profiles and nuclear separation by a separate study. Because this study focuses on generating COI sequences and documenting mitochondrial lineage structure, we retain the previously established Type A/Type B terminology and emphasize that future taxonomic work will be necessary to determine whether these lineages represent separate species. The COI sequences generated in this study provide a practical molecular tool for distinguishing these lineages and support regulatory decision-making, quarantine measures, and the development of lineage-specific pest management strategies. This will strengthen regional and national biosecurity, particularly in regions where introductions and secondary spread remain ongoing. In addition, the degree of divergence, combined with the morphological differences between Type A and Type B, falls within a range where species limits cannot be confidently inferred and therefore warrants future work.

## CONFLICT OF INTEREST

The authors have declared that they have no conflict of interest.

## AUTHORS’ CONTRIBUTIONS

**Muhammad Z. Ahmed**: study conceived, funding acquisition, conceptualization, investigation, evolutionary inference, taxonomic inference, USDA permit acquisition, coordination of collaborators and international sample transfers, methodology, software, formal analysis, visualization, data curation, project administration, supervision, writing – original draft, writing – review and editing. **Caroline Hauxwell**: sample collection, investigation, validation, writing – review and editing. **Ynna Hernandez Europa**: field sample collection, investigation, validation, writing – review and editing. **Isaac L. Esquivel**: field sample collection, investigation, data curation, validation, writing – review and editing. **David R. Kerns**: field sample collection, investigation, validation, writing – review and editing. **Nicole Quinn**: field sample collection, investigation, validation, writing – review and editing. **Darcy Patrick**: field sample collection, investigation, validation, writing – review and editing. **Sachin Rustgi**: methodology, supervision, validation, writing – review and editing. **Peilin Tan**: morphological identification, slide preparation, microscopy, slide curation, investigation, validation, data curation, writing – review and editing. **Nisha Yadav**: communication with collaborators, sample organization, DNA extraction, PCR, sequencing preparation, methodology, data curation, validation, writing – review and editing. **Blake Wilson**: field sample collection, investigation, validation, writing – review and editing.

## FUNDING INFORMATION

This work was supported by projects SC1700692 (S1073), NCERA 224, and SC1700696. Australian specimens were provided with the financial support of Meat & Livestock Australia (MLA), the Australian Government, and the Queensland University of Technology (QUT).

## DATA AVAILABILITY STATEMENT

All raw sequence data, distance matrices, R scripts, and figure files used in this study are openly available through Mendeley Data. Ahmed, Muhammad (2026), Development of the First Cytochrome Oxidase I Barcode and Evidence for a Single Haplotype Associated with the Recent United States Invasion of the Pasture Mealybug Heliococcus summervillei (Pseudococcidae, Hemiptera), Mendeley Data, V1, doi: 10.17632/7xmt6gdtx2.1.

## ACKNOWLEDGEMENTS

The authors gratefully acknowledge the students and staff at the Pee Dee Research and Education Center, Clemson University, Florence, South Carolina, USA, including Shawn Celeste Chandler and Powlomee Mondal, for their invaluable support with sample processing. We thank Muhammad Iqbal Khan (Punjab Agriculture Department, Multan, Pakistan), Ian Gibbs (Plant Protection Department, Ministry of Agriculture, Food and Nutritional Security, Government of Barbados, Barbados), and Greg Evans (USDA APHIS) for assistance with collections. Australian specimens were provided with the financial support of Meat & Livestock Australia (MLA), the Australian Government, and the Queensland University of Technology (QUT). This work was supported by projects SC1700692 (S1073), NCERA 224, and SC1700696.

## SUPPLEMENTARY MATERIALS

**Table S1.** Collection metadata for *Heliococcus summervillei* specimens used in this study, including NY codes, collection dates, host plants, geographic origin, specific locality, and collectors. The dataset includes representatives of both the historical Type A lineage (Barbados) and the invasive Type B lineage (Australia, Pakistan, U.S.). All specimens were preserved in 95% ethanol and stored at −20 °C prior to DNA extraction.

**Table S2.** GenBank COI reference sequences (*n* = 150) retained after BLAST filtering at 100% query coverage, including accession numbers, species names, and haplotype assignments.

**Table S3.** Model-selection results for all nucleotide substitution models evaluated for the COI dataset. The table reports the number of parameters, Bayesian Information Criterion (BIC), corrected Akaike Information Criterion (AICc), log-likelihood (lnL), proportion of invariant sites (I), gamma-shape parameter (G), rate-matrix values (R), and nucleotide frequencies for each model tested. Models include GTR, TN93, HKY, T92, JC, and K2 variants, each evaluated with and without gamma-distributed rate heterogeneity and invariant sites. The GTR+G model yielded the lowest BIC and AICc values and was therefore selected as the best-fit model for all ML phylogenetic reconstructions presented in Figure 1 and Supplementary Figure S1.

**Table S4.** Kimura-2-Parameter (K2P) genetic distances among COI haplotypes. Pairwise K2P distances calculated in MEGA from the trimmed 708 bp COI alignment. Distances are reported for all 1 to 58 ingroups and one outgroup 66 identity-based haplotypes defined in the 100% similarity clustering analysis. The table provides raw pairwise genetic distances used for downstream lineage comparisons (Type A, Type B), Maximum Likelihood phylogeny reconstruction, and statistical parsimony haplotype network analysis. These distance values correspond directly to the identity-based haplotype groups listed in Supplementary File S9 and Supplementary File S10.

**Figure S1.**
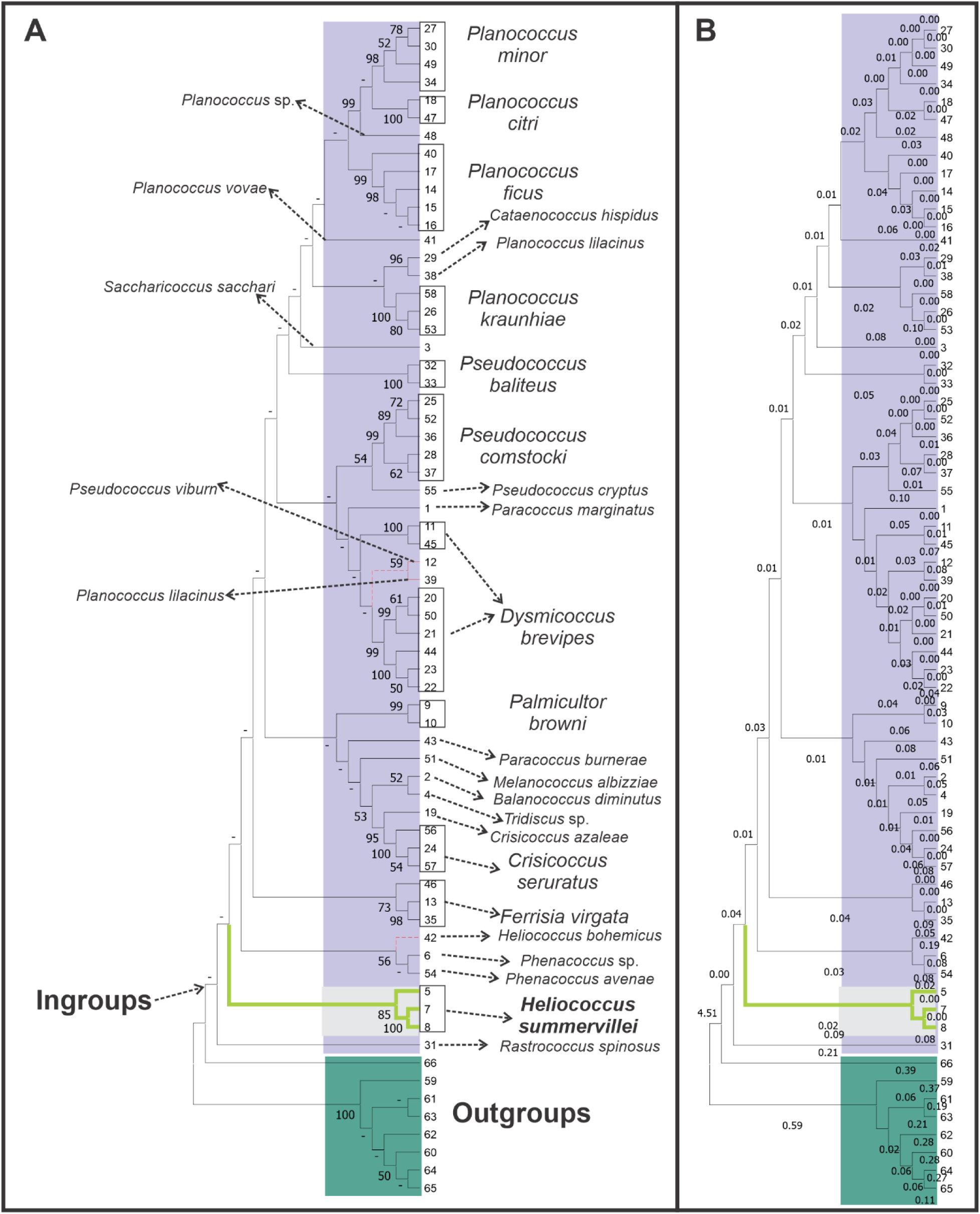
Maximum Likelihood phylogenetic reconstruction was performed under the GTR+G substitution model, identified as the best-fit model based on the lowest AICc and BIC values (Table S3). (A) Bootstrap consensus tree showing node support values. Purple shading indicates ingroup Pseudococcidae; green shading indicates non-Pseudococcidae Coccoidea outgroups; grey shading highlights *Heliococcus summervillei* Type A and Type B. Both lineages form a single clade (bootstrap = 85), and the two Type A haplotypes cluster together with full support (bootstrap = 100). Red dashed lines indicate taxa placed outside their expected clades, including haplotype 66 (non-Pseudococcidae) clustering within Pseudococcidae, two *Pseudococcus* haplotypes placed within the *Dysmicoccus brevipes* clade, and *Heliococcus bohemicus* appearing within the *Phenacoccus* clade. (B) Branch-length ML tree showing relative genetic distances among haplotypes. Type A and Type B again form a cohesive lineage within Pseudococcidae and cluster adjacent to *H. bohemicus*. Outgroups form a distinct, deeply diverged clade consistent with their placement outside Pseudococcidae. Group numbers shown in the tree correspond to identity-based clusters defined in Table S2. These numbers reflect raw pairwise distance groupings and are included here to illustrate the complete haplotype-level structure underlying the collapsed species-level representation in Figure 1. Bootstrap values < 50 were replaced with a dash (–) for clarity.

**Figure S2.**
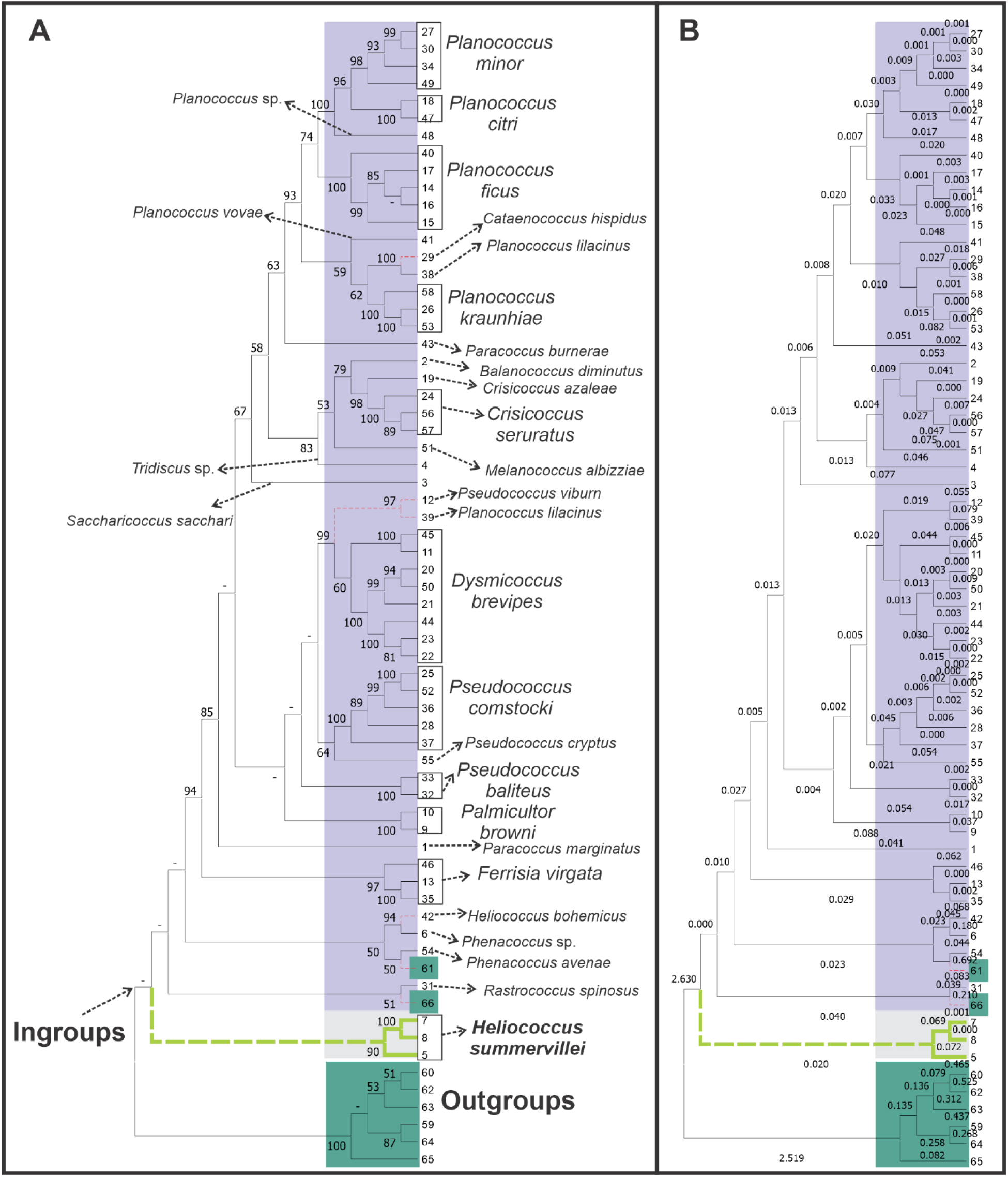
Maximum Likelihood phylogenetic reconstruction was performed in IQ-TREE v1.6.11 under the TIM+F+R5 substitution model, identified by ModelFinder as the best-fit model based on BIC (Supplementary File S8). Rate heterogeneity was modeled using the FreeRate approach with five rate categories. Purple shading indicates ingroup Pseudococcidae; green shading indicates non-Pseudococcidae outgroups (alignment positions 59–66); grey shading highlights *Heliococcus summervillei* Type A and Type B. (A) Bootstrap consensus tree showing node support values. Node support values represent a branch-length ML tree with SH-aLRT and UFBoot support values (left), and a bootstrap support tree with branch lengths scaled to genetic distances(right) percentages. Both *H. summervillei* mitochondrial lineages form a cohesive clade, with the two Type A haplotypes receiving support (90) and Type A and Type B lineages receiving full support (100). Two outgroup haplotypes (61 and 66) were recovered within the ingroup with weak support, and *H. bohemicus* was placed inside the *Phenacoccus* cluster with moderate support, although its overall placement was not well supported. Red dashed lines indicate anomalous placements relative to expected genus-level groupings. Group numbers correspond to identity-based clusters defined in Table S2. (B) Branch-length ML tree showing relative genetic distances among haplotypes. Branch lengths were optimized under the TIM+F+R5 model and illustrate the deep divergence between outgroups and Pseudococcidae. Type A and Type B again form a cohesive lineage and cluster adjacent to *H. bohemicus*. Outgroups form a distinct, deeply diverged clade consistent with their placement outside Pseudococcidae. Group numbers correspond to identity-based clusters defined in Table S2 and illustrate the underlying haplotype-level structure. Bootstrap values < 50 were replaced with a dash (–) for clarity.

## SUPPLEMENTARY FILES

**Supplementary File 1.** R-based Nuclear-mitochondrial DNA (deoxyribonucleic acid) segment (NUMT) screening workflow for newly generated COI (Cytochrome Oxidase I) sequences. This script performs sequence cleaning, gap removal, translation under the invertebrate mitochondrial genetic code (NCBI table 5), and automated detection of nuclear mitochondrial pseudogene (NUMT) signatures, including premature stop codons, frameshifts, and ambiguous amino acids (“X”). Translation confirmed correct codon phasing and absence of internal stop codons across all sequences, verifying positional homology and confirming that all COI fragments represent functional mitochondrial copies rather than NUMTs. This workflow underpins the NUMT validation results summarized in Supplementary File S2.

**Supplementary File 2.** Nuclear-mitochondrial DNA (deoxyribonucleic acid) segment (NUMT) screening results generated using the pipeline in Supplementary File S1. All newly generated COI (Cytochrome Oxidase I) sequences (*n* = 15) were translated under the invertebrate mitochondrial code and inspected for NUMT-associated disruptions. Every sequence exhibited a fully intact open reading frame (ORF), lacked frameshifts, premature stop codons, and ambiguous residues, and translated into a complete ∼230 aa COI peptide. These results confirm that all sequences represent authentic mitochondrial COI and are free of nuclear mitochondrial pseudogene contamination.

**Supplementary File 3.** FAST-All (FASTA) file containing the 15 newly generated COI (Cytochrome Oxidase I) sequences used in this study. Raw chromatograms were quality-filtered, trimmed to remove low-quality terminal bases and ambiguous characters, and aligned to ensure strict positional homology. These sequences form the core dataset for NUMT screening (Supplementary Files S1–S2), reference anchoring (Supplementary File S4), and haplotype reduction (Supplementary File S6). Metadata for each specimen is provided in Table S1.

**Supplementary File 4.** Reference-anchored alignment of all newly generated COI (Cytochrome Oxidase I) sequences against the complete mitochondrial genome of *Planococcus citri* (NCBI accession PZ166947.1). Alignment was performed in MAFFT v7 under default parameters after trimming sequences to remove low-quality terminal regions. Anchoring to a full mitogenome reference ensured that all sequences corresponded to homologous nucleotide positions within the COI gene, reconstructed a single 708-bp homologous region across samples, and prevented inclusion of truncated or mispositioned fragments. Translation under the invertebrate mitochondrial code confirmed correct codon phasing and absence of frameshifts or ambiguous residues, verifying positional homology.

**Supplementary File 5.** Comprehensive COI (Cytochrome Oxidase I) dataset comprising 165 sequences: 15 newly generated sequences (Supplementary File S3), 142 Pseudococcidae reference sequences retrieved from GenBank, and eight non-Pseudococcidae outgroups selected following the Coccoidea phylogenomic framework of Deng et al. (2025). GenBank sequences were filtered to retain only records with 100% query coverage, correct taxonomic assignment, and full positional homology to the amplified COI region. The dataset spans multiple mealybug genera (*Balanococcus*, *Cataenococcus*, *Crisicoccus*, *Dysmicoccus*, *Ferrisia*, *Palmicultor*, *Paracoccus*, *Phenacoccus*, *Planococcus*, *Pseudococcus*, *Saccharicoccus*, *Tridiscus*) and eight phylogenetically appropriate outgroups (Table S2). This alignment forms the basis for haplotype grouping (Supplementary File S6), identity-based haplotype reduction (Supplementary File S7), and Maximum Likelihood phylogenetic reconstruction (Supplementary File S8).

**Supplementary File 6.** R script and output used to collapse redundant COI (Cytochrome Oxidase I) sequences prior to haplotype analysis. This identical-sequence filtering step produced a non-redundant haplotype set used for all downstream analyses, including Maximum Likelihood phylogenetics, genetic distance estimation, and statistical parsimony network reconstruction.

Haplotype codes correspond directly to specimen identifiers in Table S1 and haplotype assignments in Table S2.

**Supplementary File 7.** Final non-redundant COI (Cytochrome Oxidase I) haplotype dataset generated after collapsing identical sequences using the workflow in Supplementary File S6. This haplotype-level alignment represents the unique haplotypes present across all newly generated and GenBank-derived sequences and was used for Maximum Likelihood phylogenetic reconstruction, haplotype network inference, and genetic distance analyses.

**Supplementary File 8.** IQ-TREE v1.6.11 output for the Maximum Likelihood phylogenetic analysis of the 708-bp COI (Cytochrome Oxidase I) haplotype alignment. ModelFinder identified TIM+F+R5 as the best-fit substitution model under BIC. The analysis included 10,000 ultrafast bootstrap replicates and SH-aLRT support values. The file contains full model selection tables, substitution matrices, FreeRate category parameters, log-likelihood scores, and both ML and consensus trees with branch lengths and support values. Four near-zero internal branches (<0.0014) were flagged for caution. This file provides the complete phylogenetic framework underlying Figures S2A–B.

**Supplementary File 9.** Analysis script used to generate the principal coordinates analysis (PCoA) and to integrate genetic distance information from Table S4 for ingroups (1 to 58) and outgroup (66). The script includes all steps required to (i) import COI (Cytochrome Oxidase I) haplotype data, (ii) incorporate pairwise K2P (Kimura 2-Parameter) distances, (iii) construct PCoA ordinations, and (iv) evaluate clustering patterns among haplotypes and lineages (Type A, Type B, excluded lineage). All functions, parameters, and workflow components needed to reproduce the PCoA and distance-based analyses presented in Figure 3A.

**Supplementary File 10.** Analysis script used to generate the lineage-structure principal coordinates analysis (PCoA) based on pairwise K2P genetic distances from Supplementary File S11 for ingroups (1 to 58). The script includes all steps required to (i) import COI (Cytochrome Oxidase I) haplotype distance data, (ii) reconstruct the full symmetric K2P matrix from the MEGA lower-triangle format, (iii) assign haplotypes to lineage categories (singleton, one-lineage species, two-lineage species), (iv) compute lineage centroids, and (v) visualize mitochondrial lineage structure using directional L1→L2 arrows. All functions, parameters, and workflow components needed to reproduce the lineage-level PCoA are provided in Supplementary Figure 3B.

**Supplementary File 11.** Pairwise Kimura-2-Parameter (K2P) genetic distance matrix derived directly from the MEGA-generated distances reported in Table S4. This file reorganizes the ingroup 1 to 58 values from Table S4 into a machine-readable matrix format suitable for downstream multivariate analyses, including Principal Coordinates Analysis (PCoA) (Supplementary File S10) and lineage-level comparisons. Distances correspond to all 58 COI haplotypes defined in the identity-based clustering analysis in Supplementary File S10.

**Supplementary File 12.** Raw graphical outputs and fully editable vector files corresponding to all figures presented in the main manuscript and supplementary materials. This archive includes the unmodified IQ-TREE phylogram (Figure 1), the branch-length phylogram (Figure 2), the Principal Coordinates Analysis (PCoA) ordination (Figure 3), and the supplementary phylogenetic figures (Figures S1–S2). For each figure, both the original analytical output (PNG) and the post-processed vector-editing files [CorelDRAW (.cdr)] are provided. These editable versions contain all graphical refinements applied during manuscript preparation, including color-coding of ingroup and outgroup taxa, annotation of bootstrap values, haplotype labels, lineage markers, and adjustments to layout, typography, and panel spacing. All edits were performed using non-destructive vector workflows to preserve full reproducibility and allow downstream modification. Together, these materials provide complete transparency for figure generation and ensure that all phylogenetic and multivariate visualizations can be independently verified, re-edited, or re-exported for future analyses.

## Notes

### Competing Interest Statement

The authors have declared no competing interest.

### Summary of Updates

This revised version of the manuscript incorporates extensive updates based on the detailed editorial assessment and referee comments received during review at the Journal of Applied Entomology. We rebuilt the dataset and analyses from the foundation upward to ensure full accuracy, positional homology, and consistency across all COI sequences. First, we corrected the marker description throughout the manuscript. The amplified fragment corresponds to the Jerry Pat region of COI, not the COI 5p barcode region. All text, methods, and figure legends have been updated to reflect this correction. We also clarified the rationale for using the Jerry Pat fragment, which is the most reliable COI region for mealybugs and is widely used in diagnostic reference libraries. Second, we reconstructed the full alignment using homologous COI sequence regions anchored to the Planococcus citri mitochondrial genome. All sequences were trimmed to a single verified region, translated under the invertebrate mitochondrial code, and checked for uninterrupted open reading frames to confirm positional homology and exclude NUMTs. Third, we corrected all genetic distance calculations. Several outgroup sequences in the earlier version were short, non overlapping, or showed NUMT like characteristics, which produced inflated K2P distances. These records were removed. Distances were recomputed using only validated sequences aligned to the same COI region. The divergence between the historical Type A lineage and the invasive Type B lineage is now consistently recovered at approximately ten percent. Fourth, we corrected all lineage assignments, identity groups, and sampling interpretations. Barbados contains one verified Type A haplotype. The invasive lineage contains one verified Type B haplotype. NY77 was retained in the dataset as a divergent specimen, but its placement is now described carefully and without over interpretation. We removed any language that overstated species boundaries or implied conspecificity. Fifth, we revised the Results, Discussion, and all comparative sections to ensure taxonomic neutrality. The manuscript now presents a cautious interpretation consistent with the corrected genetic distances, phylogenetic structure, and morphological evidence. Finally, we updated all references, corrected GenBank accession information, verified all citations, and replaced all figures and supplementary files with validated versions. This revised manuscript provides a corrected and reliable COI dataset for Heliococcus summervillei and a clear foundation for future diagnostic and taxonomic work.

